# NMR Line Shapes in Molecular Mechanisms with Ligand Binding and Multiple Conformers

**DOI:** 10.1101/2020.01.03.894485

**Authors:** Evgenii L Kovrigin

## Abstract

Interactions of ligands with biological macromolecules are sensitively detected through changes of chemical shifts and line shapes of the NMR signals. This paper reports a mathematical analysis and simulations of NMR line shapes expected in titrations when ligand binding is coupled to multiple isomerization transitions. Such molecular mechanisms may correspond to ligand binding by intrinsically disordered proteins or by autoinhibited enzymes. Based on the simulation results, we anticipate several specific effects that may be observed in practice. First, the presence of non-binding conformers of the receptor molecule leads to a remarkable broadening in the binding transition even if the exchange between binding and non-binding conformers is very slow. Second, the ligand-binding mechanisms involving induced fit are expected to demonstrate deceptively decelerated exchange regimes even when the underlying kinetics are very fast. Conversely, the observation of fast-exchange shifting resonances with modest line-broadening (“marching peaks”) in practical NMR titrations may involve conformational selection transitions but less likely to be observed for the induced fit. Finally, in auto-inhibited molecules that open to form multiple binding-competent conformers, the fast dynamics of opening/closing transition are capable of masking the true kinetics of interconversion among transiently open forms of the receptor.

## Introduction

A variety of biological functions involve ligand binding coupled to conformational changes in biological macromolecules. While folded conformations are typically unique and well defined, the “open” or unfolded forms are represented by ensembles of multiple alternative conformers. Intrinsically disordered proteins (IDP) are one example where multiple unfolded or partially folded polypeptide conformations are in exchange, and the ligand interacts with a matching one—often followed by an additional induced-fit transition to the tightly bound state^1-3^. Another situation with multiple alternative conformations arises when a protein structure undergoes a conformational transition (“opening”; local unfolding) to allow for the ligand binding, and the ligand will induce a new stable conformation. In this scenario, multiple alternative first-encounter complexes may be formed between partially structured binding site and the ligand, which then evolve into a single well-packed tightly bound conformation^4,5, 6-9^. The line shapes and the chemical shifts of NMR signals are sensitive parameters reflecting fine details of the molecular interactions and conformational transitions^10-15^. The NMR line shape evolution had been explored previously for conformational selection and induced fit^16-21^. In this study, I expanded the NMR line shape analysis to systems with multiple binding or non-binding conformations in a dynamic exchange (as observed with intrinsically disordered or autoinhibited macromolecules). Three generalized models of conformational selection and induced fit with up to five isomers were developed and implemented in MATLAB as a part of *Integrative Data Analysis Platform (IDAP).* The IDAP code was interfaced to TITAN^20^ for the two-dimensional line shape simulations.

### Theory

In what follows, the receptor, R, will be the species that is labeled with magnetically active isotope thus giving rise to the NMR signal. The ligand, L, is its binding partner and it is unlabeled—not observed in NMR spectrum. Other than that, there are no other conditions on what R and L may be (protein, RNA, oligosaccharide, small molecule, etc.).

In this paper, we will analyze two distinct situations that may be encountered in biomolecular systems with ligand binding: (1) multiplicity of the *non-binding* conformers (or isomers) of the receptor, and (2) multiplicity of *binding-competent* conformers. The first situation occurs when the receptor molecule is conformationally dynamic, and the ligand-binding site is only formed in *one* specific isomer of the R molecule, while the rest of isomers are incapable of binding the ligand. Figure 1 shows a model of this kind with five non-binding isomers of the receptor. This model is referred here to as *U-5R* following the *LineShapeKin Simulation* conventions^19^ (“*U*” stands for unimolecular ligand binding event, “*R*” indicates that the receptor is undergoing isomerization, “5” denotes five binding-incompetent conformers present alongside with a single binding-competent form). The *U-5R* model may be extended to a situation when the ligand binding to the R species results in a first encounter complex, which further isomerizes to a tightly bound form—the *U-5R-RL* model (Figure 2).

**Figure 1.**
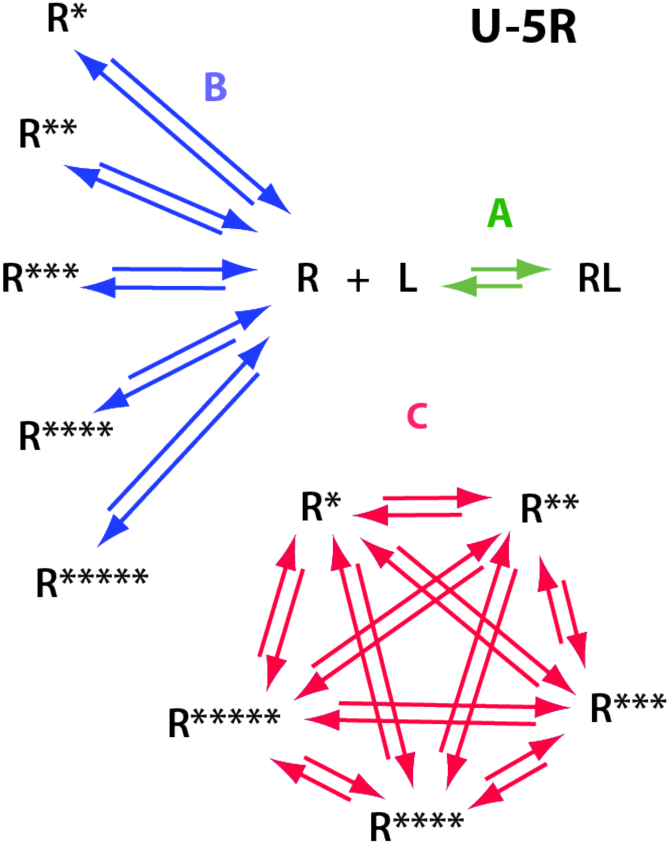
U-5R model: the receptor molecule has multiple conformations (R*, R**, R***, R****, and R*****) that are incapable of binding the ligand and are in exchange with each other (C transitions). The single binding-competent receptor conformation, R, is in exchange with all binding-incompetent species through the B transitions. The R binds the ligand, L, to form the bound species, RL (A transition). This generalized model may be adjusted to account for specific practical cases by setting necessary equilibrium and rate constants to desired values or to a zero thus eliminating specific species or transitions altogether.

**Figure 2.**
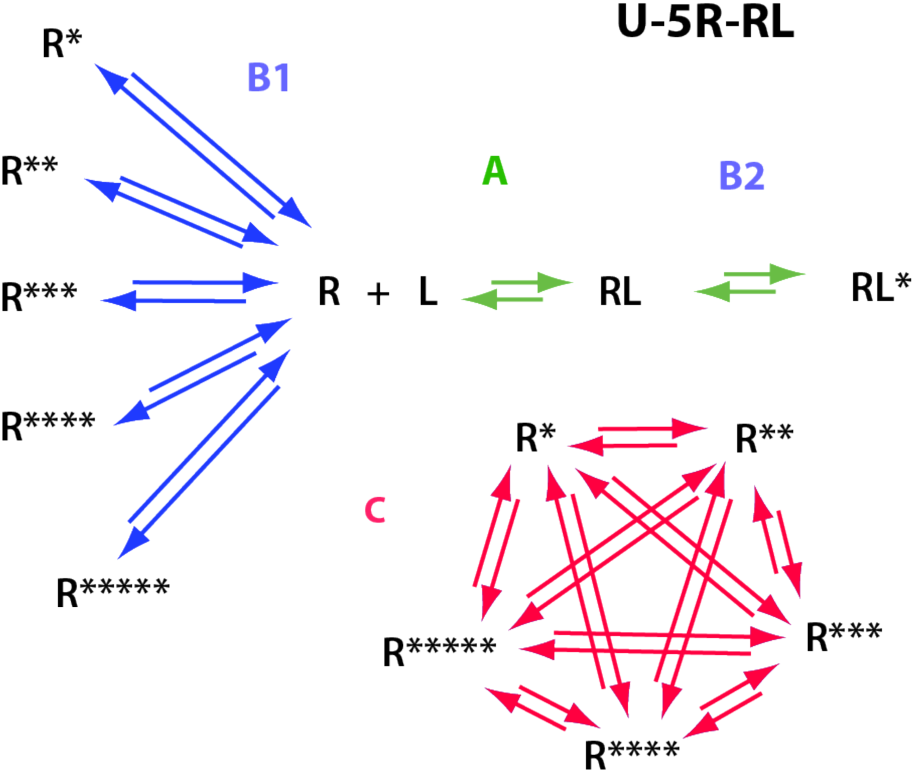
U-5R-RL model: the same as U-5R but the first-encounter receptor-ligand complex, RL, additionally isomerizes through the induced-fit mechanism to a tightly bound state, RL* (B2 transition).

An entirely different molecular mechanism operates when a receptor molecule exists in a binding-incompetent ground state with a transient formation of conformers capable of binding the ligand (Figure 3). The binding-incompetent receptor conformations like the R* in Figure 3 are commonly referred to in enzymology as *resting* or *autoinhibited* states with restricted access to their binding sites. The binding-competent isomers may form due to the conformational opening of flexible structures or through the local or global unfolding.

**Figure 3.**
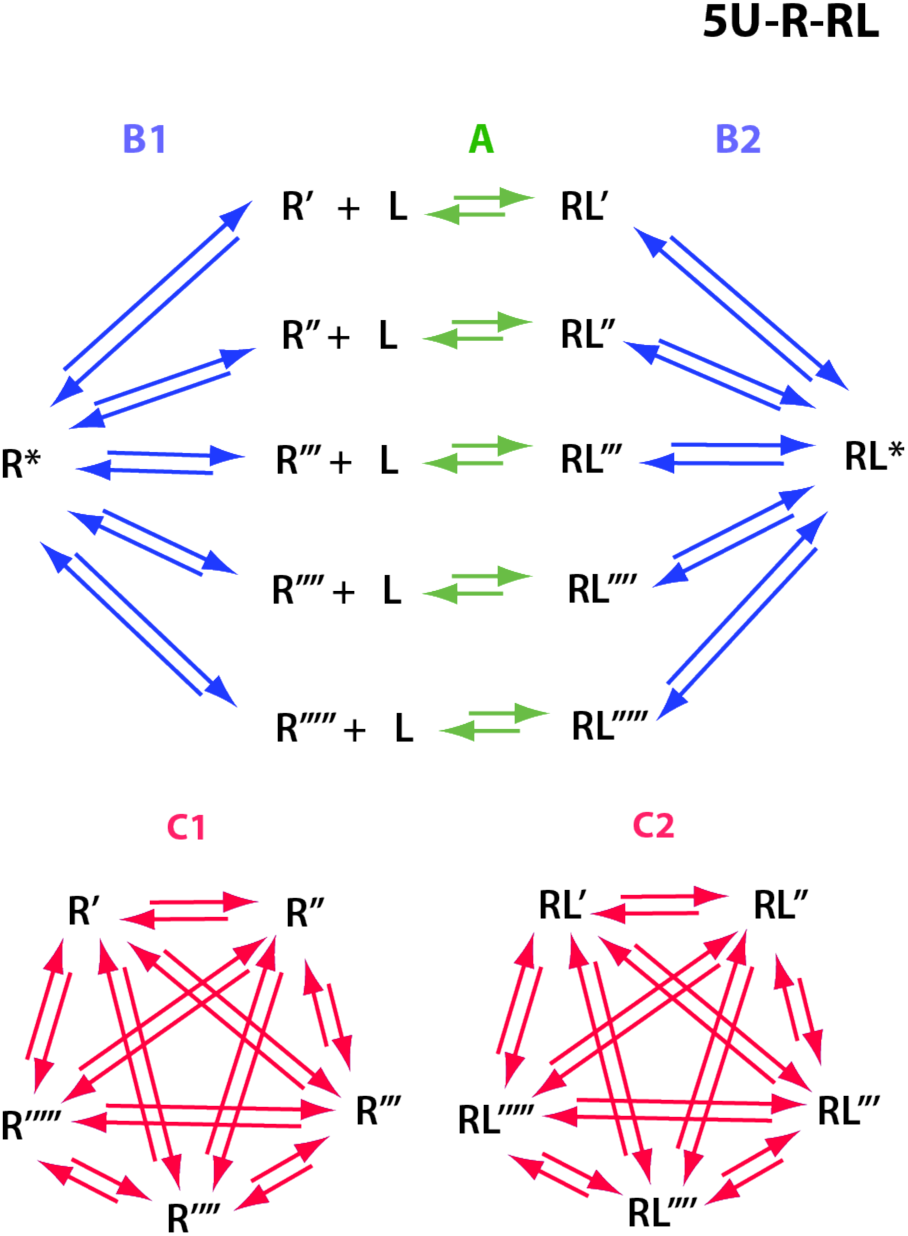
5U-R-RL model: the ground state of the receptor molecule, R*, is incompetent of binding a ligand until it isomerizes into one of the binding-competent conformations, R’ to R’’’’’ (B1 transition). The binding-competent conformations may be in exchange (C1 transitions). Multiple possible first-encounter complexes are formed upon contact with the ligand, L (A transitions), which may also be all in dynamic exchange (C2 transitions). The tightly bound species is formed in the final induced-fit step (B2 transition).

For all three models, I developed the variants with 1, 2, 3, 4, and 5 isomers. Five isomers were considered to be a large enough number to provide a good approximation of the model behaviors for all numbers of isomers beyond five. The systems of equilibrium thermodynamic equations for the coupled ligand-binding and isomerization equilibria were explicitly solved using individual equilibrium constants *for each transition*. The kinetic matrices for the NMR line shape simulations were also developed including individual kinetic rate constants for transitions between *each pair of species*.

The molecular systems with many isomerization transitions present a daunting variety of scenarios one may need to consider. Here, for simplicity, we will assume that all alternative isomers are *identical*, therefore, their formation is governed by the same equilibrium and kinetic rate constants. Identity of the isomers also means that they are equally populated (meaning unity pairwise equilibrium constants), and that all pairwise isomer exchange transitions are kinetically identical as well.

The derivations for *U-5R, U-5R-RL*, and *5U-R-RL* models were performed in MuPad (Mathworks) and implemented as a part of the model library in IDAP (Integrative Data Analysis Platform, http://lineshapekin.net/#IDAP). In all derivations, the forward and reverse kinetic rate constants were denoted as k_1_ and k_2_, respectively, and equilibrium constants in binding steps were association constants, K_a_. For clarity of presentation in the figures, k_2_ of ligand-binding steps was labeled as k_off_ and K_a_ was converted to the dissociation constants, K_d_. For simulations of the two-dimensional HSQC NMR spectra, the IDAP models were interfaced to TITAN^20^. The MuPad notebooks detailing the thermodynamic and kinetics derivations as well as the MATLAB code of the IDAP are available from the author upon request. Full details of simulations for the figures in the manuscript are provided in the Supporting Information. TITAN may be obtained from https://www.ucl.ac.uk/biosciences/christodou-lou-group/research/titan-2d-nmr-lineshape-analysis. All NMR spectra were simulated assuming a static magnetic field of 14.1 T.

## Results and Discussion

Multiplicity of alternative transitions creates a universe of possible exchange scenarios because the coupled transitions may be in the fast, intermediate, or slow exchange regimes (in any combinations). In what follows, we will not attempt to cover this overwhelming variety but, instead, highlight some remarkable phenomena to stimulate reader’s interest in this topic.

### Reference simulation

We will start with a simple 1:1 binding model to create a “baseline” view of the NMR titration. In the next steps, we will expand it into the mechanisms with multiple conformational selection and induced fit transitions. Figure 4 shows a simulated titration series for a modest K_d_ of 10^−5^ and the off-rate constant of 3000 s^-1^, typical of a weak-binding ligand.

**Figure 4.**
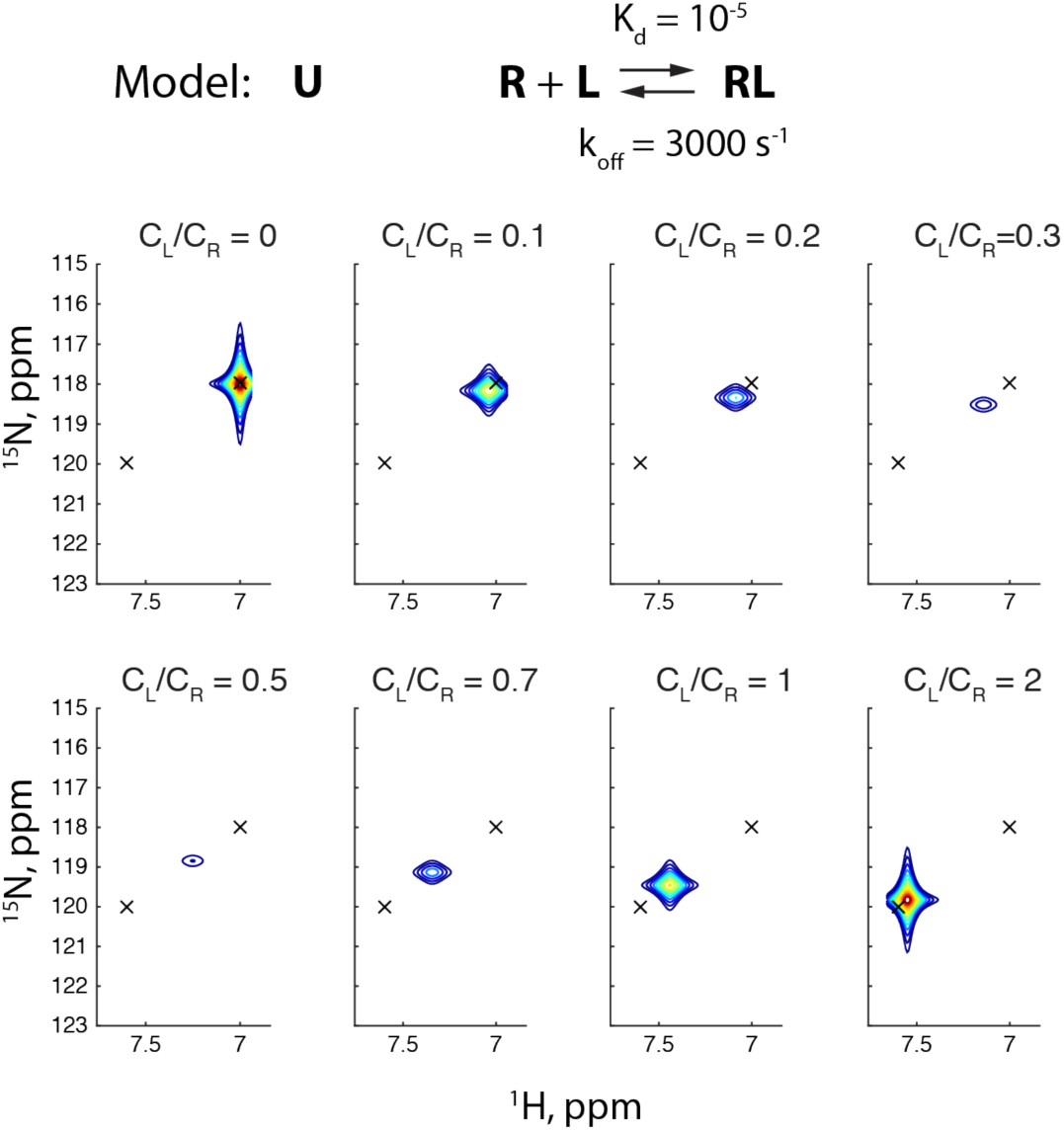
A reference simulation for the 1:1 binding mechanism (lock-and-key; U model) in fast-intermediate exchange. Two-dimensional ^1^H-^15^N HSQC spectra were calculated using TITAN^20^. The frequency differences between R and RL states are 2261 s^-1^ in ^1^H, and 754 s^-1^ in ^15^N dimensions, respectively. Molar ratios of ligand to receptor in the titration are indicated above each spectrum. The lowest contour is set at 1% of the maximum peak height in the titration series with a factor of 1.4 for the following contours. For full details of the simulation setup see *simulations/U_1R_simulated_U*.

As it might be anticipated for the chosen kinetic parameters and frequency separations in Figure 4, the titration series features a weighted average peak that gradually shifts from the position of the unbound receptor (at C_L_/C_R_=0) toward the position of the receptor-ligand complex (C_L_/C_R_=2). The peak is significantly broadened in proton dimension in the intermediate titration points (C_L_/C_R_=0.5) corresponding to the fact that the binding off-rate constant only slightly exceeds the frequency separation of species in the proton dimension (k_off_=3000 s^-1^ and Δ*ω*=2261 s^-1^, respectively) resulting in the fast-intermediate exchange regime (for definition of the exchange regimes see Kaplan and Fraenkel^22^). The nitrogen dimension shows less peak broadening because of a smaller frequency difference between the exchanging sites, Δ*ω*=754 s^-1^. This simple titration series will help us spot unintuitive features of the mechanisms with multiple alternative isomers.

### Simple conformational selection

In the next step, we add an isomer of a receptor that is incompetent for ligand binding, R*, producing the U-1R model (Figure 5). We will set the exchange kinetics between R and R* to a very slow exchange regime by choosing the rate constants for the R ↔ R* isomerization equal to 1 s^-1^, for clarity of observations. Correspondingly, the HSQC spectrum in the absence of the ligand, at C_L_/C_R_=0 in Figure 5, shows two peaks with identical intensities. Addition of a ligand to 10% saturation (C_L_/C_R_=0.1) leads to broadening and shifting of the R peak. If compared to the U model titration, the R peak evolution resembles the line shapes of the simple 1:1 binding model while the R* gradually diminishes in-place. In this slow-exchange regime, the two transitions in the molecular mechanism appear uncoupled due to a large difference in the exchange rates. Yet, peak broadening of this model with a conformational selection step is more significant in Figure 5 in the *early titration points* (at C_L_/C_R_=0.1-0.3) than in Figure 4 while the opposite is true in the later stages of titration.

**Figure 5.**
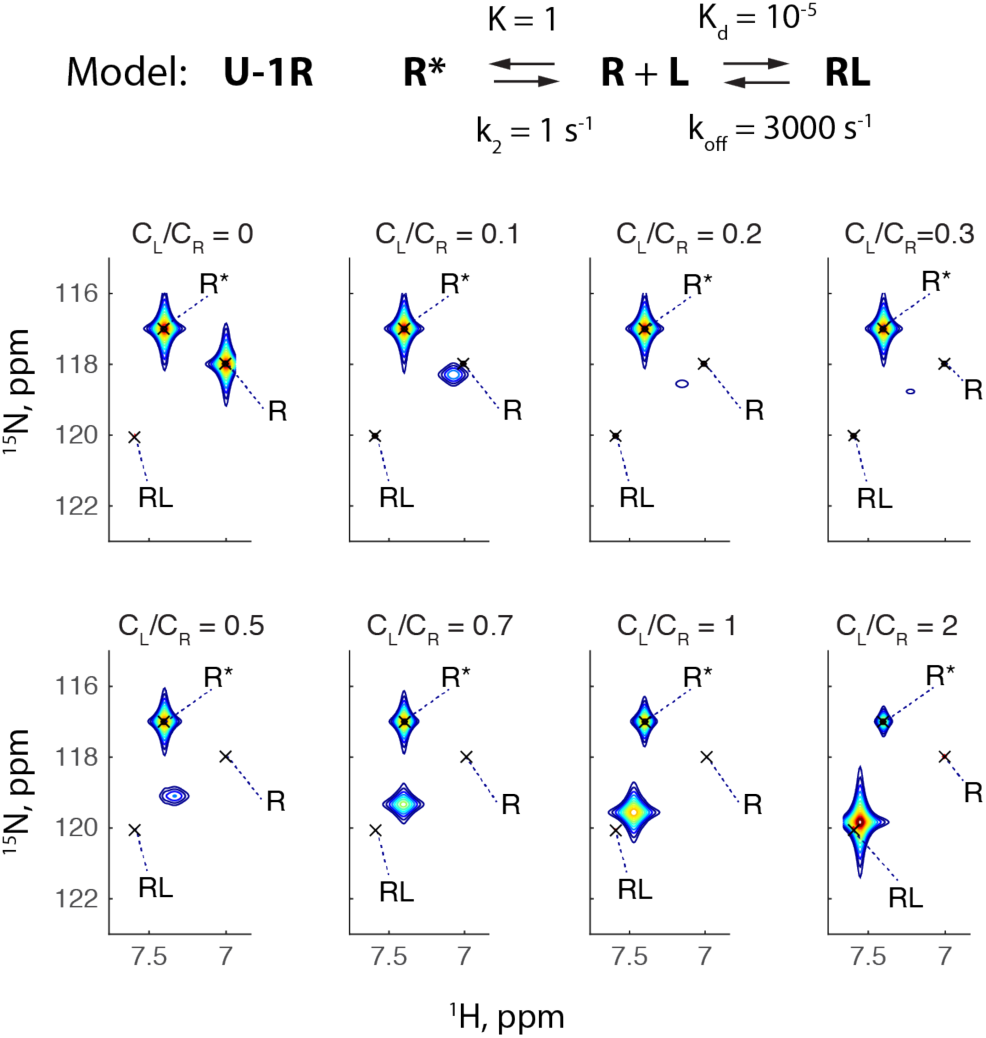
2D ^1^H-^15^N HSQC NMR spectra for the U-1R model. The titration schedule is the same as in Figure 4. Chemical shifts of species (^1^H, ppm/^15^N, ppm): R, 7.0/118; R*, 7.4/117; RL, 7.6./120. The frequency differences between R and R* states are 1507 s^-1^ in ^1^H and 377 s^-1^ in ^15^N dimensions, respectively. The rate constant k_2_ is a reverse rate constant and K is an equilibrium constant for R* formation from R. Contour levels are the same as in Figure 4. Populations of species are shown in Supporting Figure 1. For full details of the simulation setup see *simulations/U_1R_Bslow_Afast_3*.

### Conformational selection in slow-intermediate exchange

If we accelerate the exchange regime in the R-to-R* isomerization transition to 80 s^-1^ (which is still in slow exchange) we clearly see how the coupled transitions interact (Figure 6). Broadening of the weighted-average peak in the early titration points is dramatically increased: the R peak becomes, practically, undetectable upon addition of the very first ligand aliquot (C_L_/C_R_=0.1). The weighted-average resonance re-emerges at as late as 50% saturation (C_L_/C_R_=0.5). Thus, the slow-intermediate exchange dynamics in the conformational selection step dramatically increases the exchange broadening in the binding reaction. It is notable that the binding reaction is expected to follow the fast-exchange pattern as it starts with k_ex_ of 3000 s^-1^, which is greater than the frequency difference Δ*ω*=2261 s^-1^. The expected fast-exchange regime should become even faster as more ligand is added because the k_ex_ increases according to the basic relationship *k*_*ex*_ = *k*_*off*_ + *k*_*on*_[*L*]. Yet the observed exchange regime appears to be intermediate.

**Figure 6.**
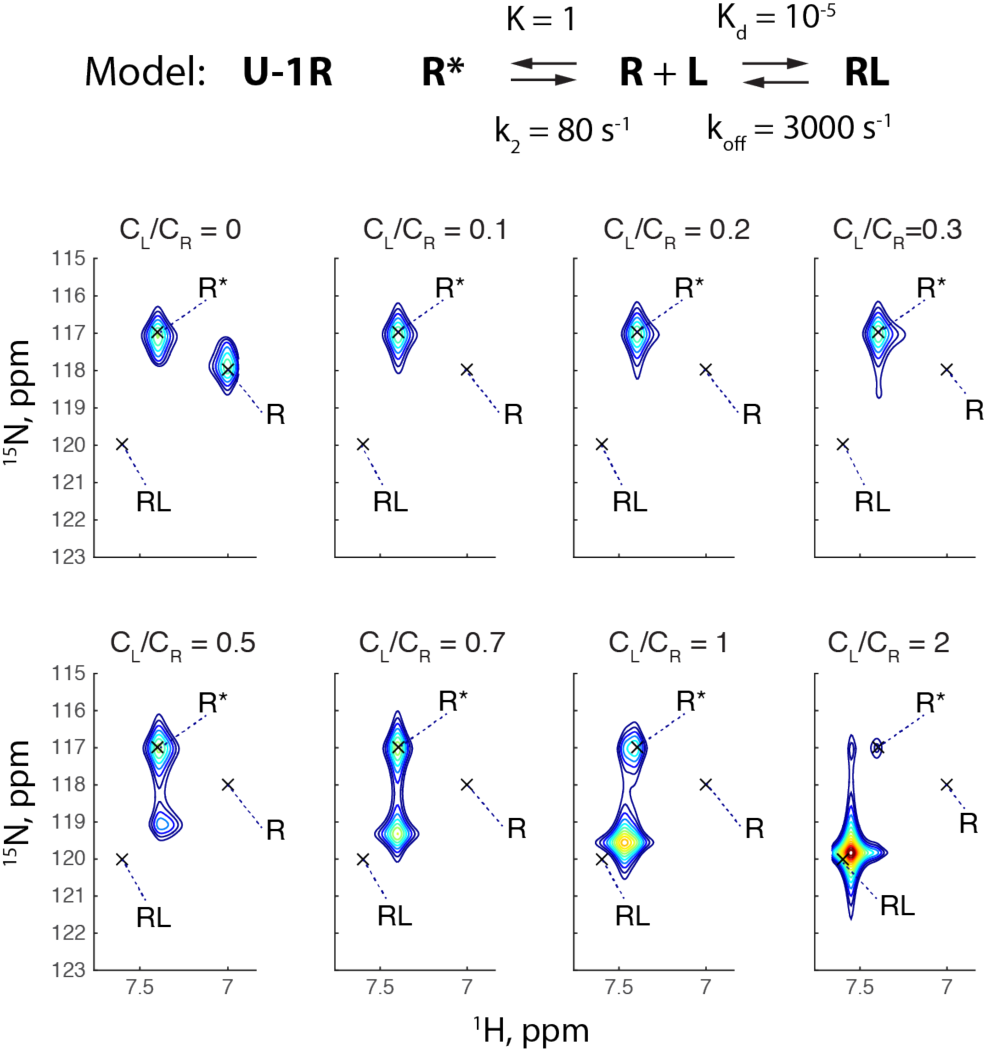
2D ^1^H-^15^N HSQC NMR spectra for the U-1R model with the rate constant for formation of R* set to 80 s^-1^. Contour levels and ppm positions of all species are the same as in Figure 5. Populations of species are shown in Supporting Figure 1. For full details of the simulation setup see *simulations/U_1R_Bint_Afast_3*.

### Exchange peaks in 2D simulations

It is notable that the simulations of the two-dimensional line shapes in Figure 6 feature the exchange cross-peaks such as the one seen for the weighted-average peak and the R* peak in the HSQC for C_L_/C_R_=2. Appearance of these peaks is well anticipated as the HSQC experiment contains INEPT coherence transfer periods during which the exchange mixing occurs to a certain degree. The TITAN algorithm directly simulates the HSQC pulse sequence^20^ thus accounting for the exchange during INEPT periods. Waudby and co-workers thoroughly analyzed this phenomenon in a recent publication^23^ using a classical example of conformational exchange, N,N-dimethyltrichloroacetamide, DMTCA ^24^. This small molecule has a hindered rotation with 50% populations of two states. Equal populations create most favorable conditions for exchange as manifested by large exchange cross-peaks in the longitudinal exchange experiment (see Figure 1 in Waudby et al., 2019^23^).

One important aspect of the exchange peaks in Figure 6 is that their intensity is calculated to be quite weak—only two contour levels, which corresponds to 1.4% of maximum spectral intensity in the spectral series. Our simulation is noiseless; this is why the exchange peak is observable near the R* peak in the HSQC for C_L_/C_R_=2 (at 7.55 ppm ^1^H and 117 ppm ^15^N), yet its counterpart near 120 ppm ^15^N is not—overlapped with the shoulder of a weighted-average resonance of the binding transition.

Obviously, observation of a peak at about 1% height of the main resonance needs the signal-to-noise ratio greatly exceeding 100, which is rare in biomolecular experiments. Yet, direct line shape analysis of such spectra may be informative in studies of small molecules where high sensitivities are easily achievable with the use of ultra-high magnetic fields and cryogenically cooled probes^23^.

### Two non-binding conformers in slow exchange

Our next step will be to add a second non-binding isomer, R**, to the U-1R model thus making the U-2R model (Figure 7). Remarkably, the presence of R** dramatically increases exchange broadening in the HSQC spectra in all titration points. It is important to note that the TITAN algorithm explicitly simulates the signal loss during the magnetization transfer steps of the pulse sequence due to transverse relaxation^20^. Therefore, any exchange-broadened resonance will have relatively smaller volume in the simulated spectrum than otherwise would be expected from the population of the corresponding species. Indeed, the spectrum for the receptor in the absence of a ligand, C_L_/C_R_=0, in Figure 7 demonstrates that while the populations of the R, R*, and R** forms are equal, the R and R* peaks appear significantly weaker than R**. The R and R* peaks are closer to each other than to R**, therefore, they suffer increased slow-intermediate exchange broadening relative to the more distant R** resonance. Overall analysis of the peak volume in Figure 7 shows that if the total peak volume of the final titration point (C_L_/C_R_=2) is set to 100%, then all peaks in the first titrations point add up only to 40% (at C_L_/C_R_ =0) while the sixth point (at C_L_/C_R_ =0.7) has only 52% of the final volume. This observation reflects a known difficulty of experimental work with the exchanging systems—they demonstrate reduced signal-to-noise ratios even if peaks are spectrally resolved.

**Figure 7.**
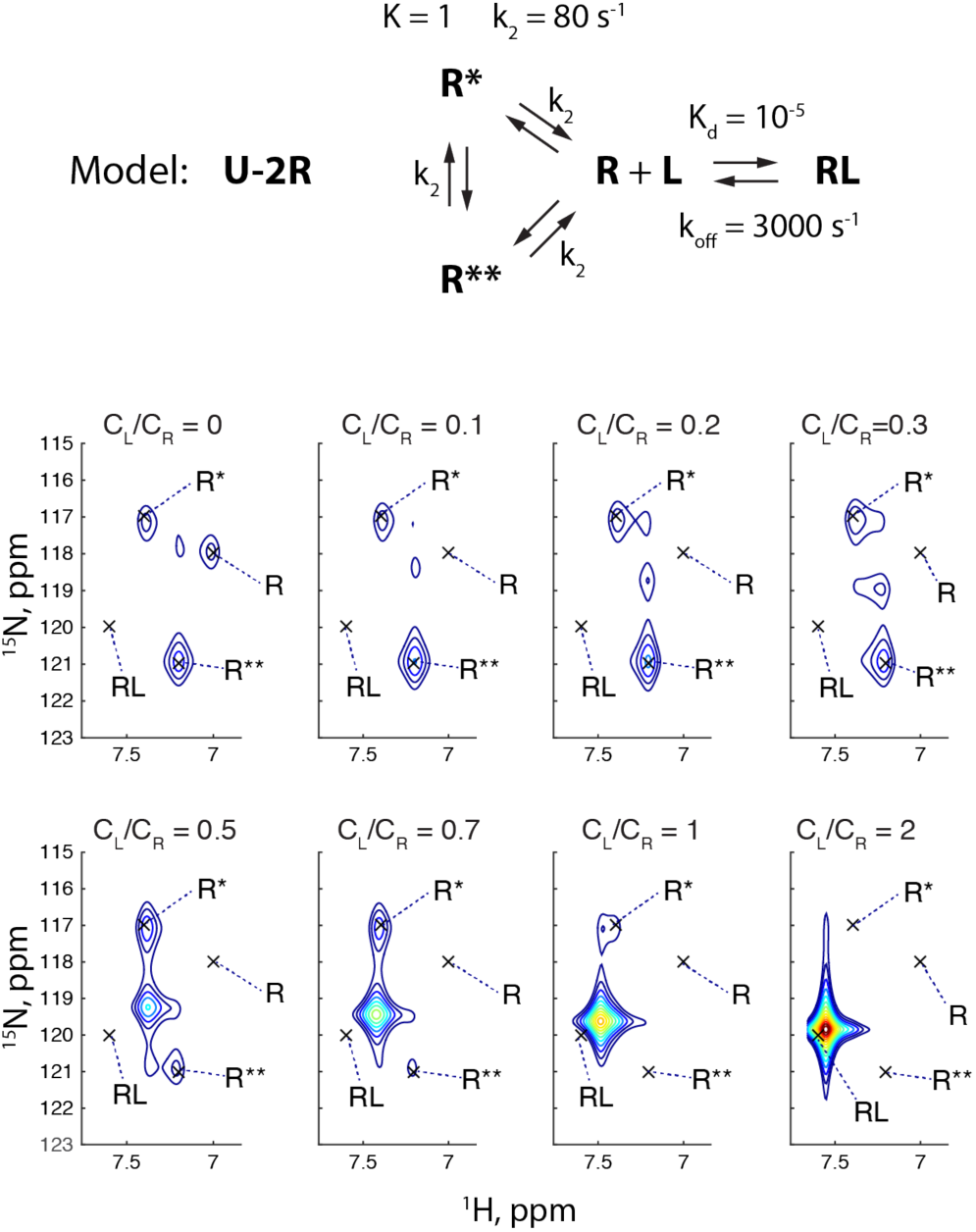
2D ^1^H-^15^N HSQC NMR spectra for the U-2R model. Chemical shifts of the R** species were set to 7.2 ppm and 121 ppm in proton and nitrogen dimensions, respectively. Populations of species are shown in Supporting Figure 2. For full details of the simulation setup see *simulations/ U_2R_Bint_Afast_3*.

### Transient narrowing of slow- and fast-exchange peaks

Another remarkable phenomenon in Figure 7 occurs when the weighted average peak of the ligand-binding transition shifts and crosses the proton frequencies of R** and, later, R* species (at C_L_/C_R_ = 0.2 and 0.5). At these titration points, the corresponding slow exchange resonances of pure R** and R* species intensify (compare numbers of contours in the R** and R* peaks at C_L_/C_R_ = 0, 0.2, and 0.5). This phenomenon resembles the transient narrowing described earlier for the three-state mechanisms in fast exchange^19^. The origin of the transient peak narrowing is that the weighted-average resonance of the binding transition may be treated as a pseudo-species in exchange with the non-binding isomers R* and R**. As the frequency of this pseudo-species shifts and matches the frequencies of one of the isomers, the Δ*ω* vanishes and the exchange broadening reduces thus making peaks narrower.

In my previous report, the transient narrowing condition was described in the fast-exchange regime in one-dimensional simulations^19^. Utilizing TITAN, this effect may be readily visualized in two dimensions for any of the multi-state models. In Figure 8, I simulated the full U-5R model (Figure 1) involving five non-binding conformers in exchange with the binding-competent receptor form, R. The transient narrowing condition requires chemical shifts of non-binding species to lie in between ligand-free and lig- and-bound receptor states (marked as *iso* in the Figure 8). In Figure 8, the exchange regime among isomers was set to fast intermediate resulting in a single weighted-average resonance shifting in the course of titration. We observe that the peak intensity in the third titration point (at C_L_/C_R_ = 0.2), as the peak crosses the “field” of the non-binding conformers, becomes greater than in the initial (ligand-free, C_L_/C_R_ = 0) or the following (C_L_/C_R_ = 0.5, 0.7, and 1.0) titration points (the greater number of red contours indicates greater intensity). Supporting Figure 4 shows this titration series in a more familiar one-dimensional representation.

**Figure 8.**
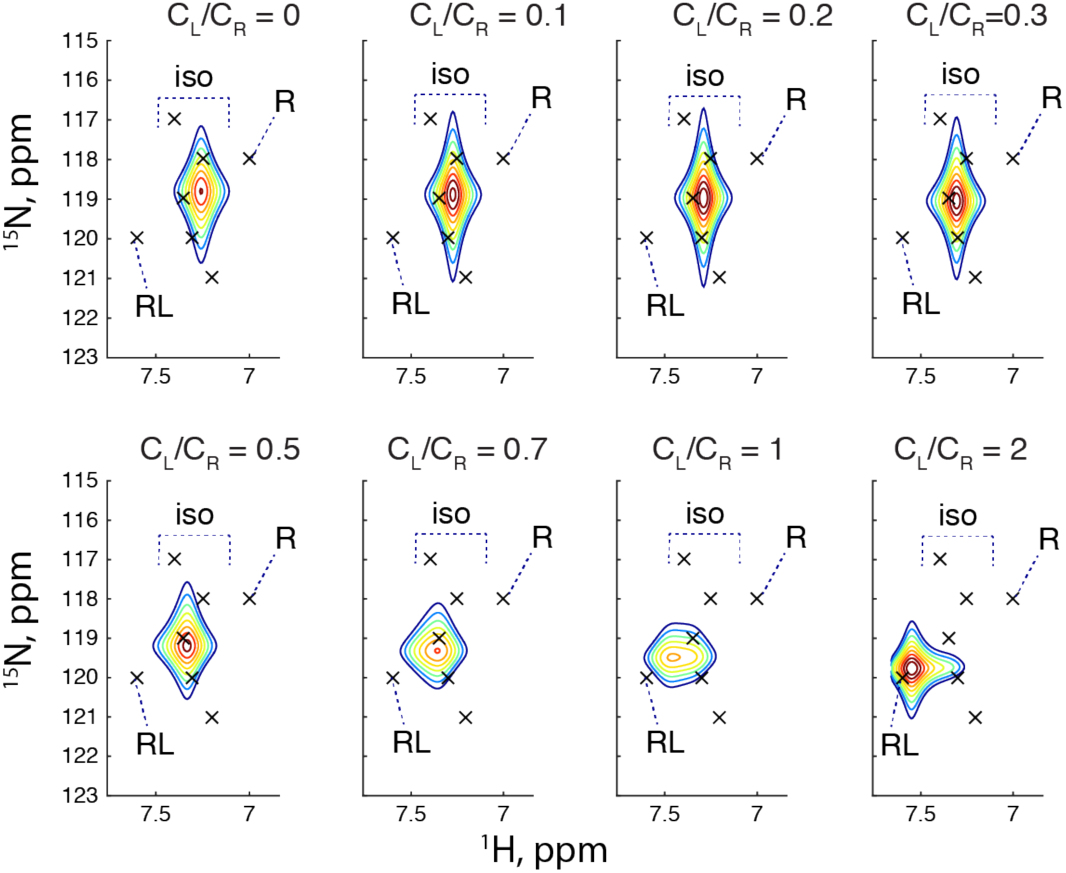
Transient narrowing in the U-5R model. 2D ^1^H-^15^N HSQC NMR spectra for the U-5R model were simulated with the reverse rate constants for formation of R isomers (k_2,B_ and k_2,C_) set to 300 s^-1^. Populations of all un-bound receptor forms (R, R*, R**, R***, R****, and R*****) are equal. Binding step has the reverse rate constant of 3000 s^-1^. Titration schedule is the same as in previous simulations. Populations of species in a titration are shown in Supporting Figure 3. Molar ratio of ligand over receptor is given in each 2D HSQC header in the top panel. Spectral positions of the non-binding conformers (R* to R*****) are labeled with ‘iso’. For full details of the simulation setup see *simulations/ U_5R_transient_narrowing*.

It notable that the transient narrowing (as in the Figure 8) was previously demonstrated for the fast-exchange regime^19^ but not for the *slow-exchange* resonances (as seen in Figure 7). Two-dimensional simulations are required for analysis of such slow-exchange peaks in the presence of fast-exchange coupled transitions. Re-examination of Figure 7 reveals that the transient narrowing only occurs when the moving weighted-average peak crosses the proton frequency of the stationary slow-exchange resonance (there is no crossing in the nitrogen dimension with the current choice of chemical shifts). When such proton frequency crossing occurs, the slow-exchange peak becomes more intense and narrow but the peaks now are completely overlapped in the proton dimension—the condition that precludes observation of individual line widths by the one-dimensional simulations (and one-dimensional NMR experiments, in general).

#### Induced-fit step and unanticipated deceleration of the exchange regime

The U-5R model shown in Figure 1 may represent a binding mechanism of an intrinsically disordered protein with just one of the conformers forming the binding surface with the ligand. Such interactions might often result not in a final bound state but a transient first-encounter complex that further isomerizes into a tightly bound species. This situation is described by the U-5R-RL model (Figure 2), which expands the U-5R model through an addition of the induced-fit step.

To appreciate the effect of the induced fit on the line shapes we will, first, examine a titration for the U-5R model in Figure 9. For this simulation I set *all* steps of the mechanism to a fast-exchange regime: k_ex_ values are in excess of 3000 s^-1^ for all transitions, which is greater than the 800-1500 s^-1^ range of frequency differences between exchanging species in Figure 9. Panel A shows the U-5R simulation producing a familiar “marching peak” pattern expected from the fast exchange condition *k*_*ex*_ > Δ*ω*.

**Figure 9.**
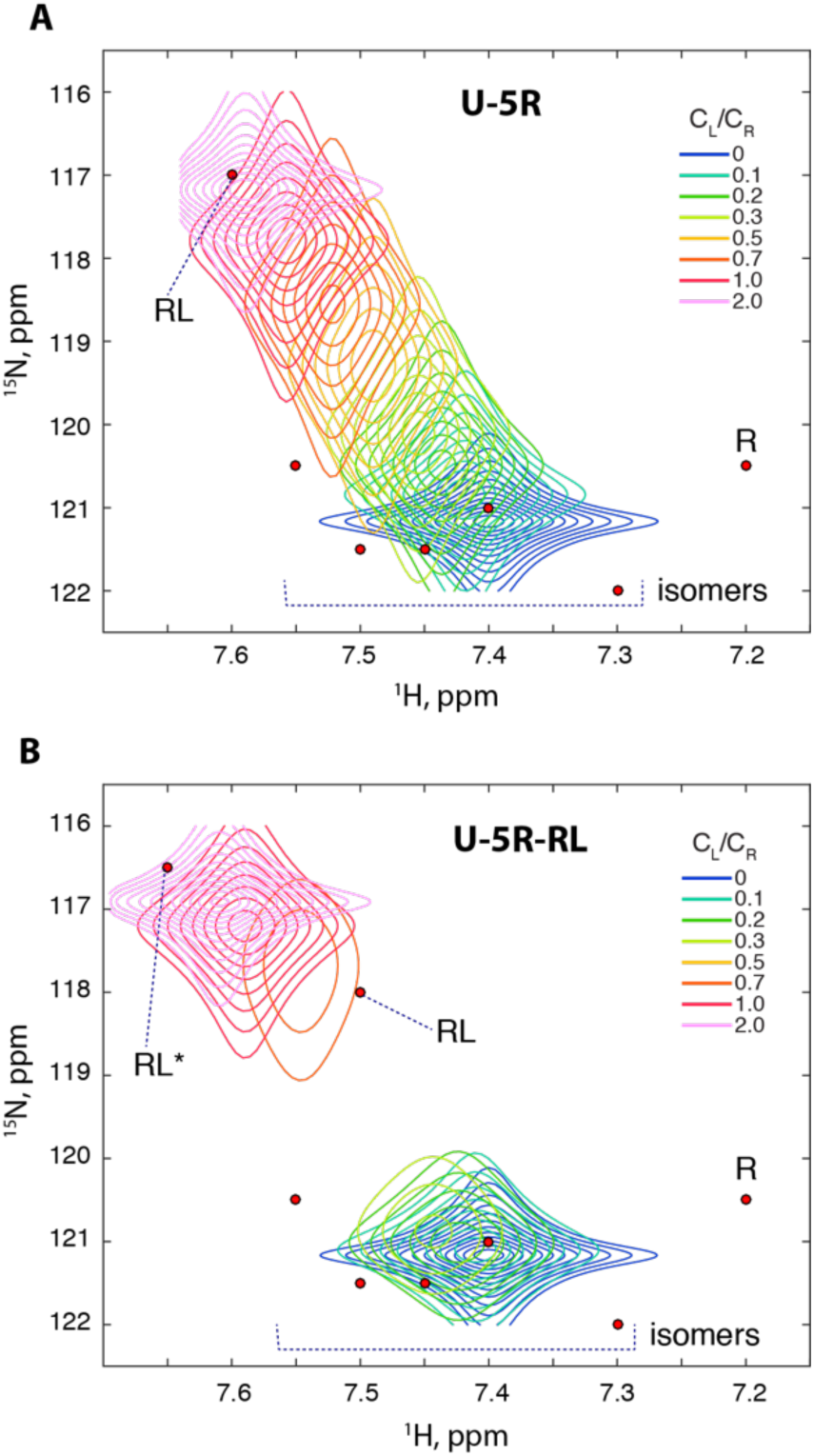
2D ^1^H-^15^N HSQC NMR spectra simulated for U-5R (**A**), and and U-5R-RL (**B**) models with all kinetic constants (k_2,A_, k_2,B_ and k_2,C_) set 3000 s^-1^. Populations of all unbound receptor forms (R, R*, R**, R***, R****, and R*****) are the same and equal to 17%. Titration schedule is the same as in previous simulations. Populations of species are shown in Supporting Figure 5. The chemical shifts of RL and RL* in **B** were adjusted to have the weighted-average peaks start and finish at the same positions in the spectral planes in both panels. For full details of the simulation setup see *simulations/U_5R* and *simulations/U_5R_RL*.

Panel B in Figure 9 shows a simulation of the U-5R-RL model where the induced-fit transition was added to the binding reaction. For visual clarity and ease of comparisons, the frequencies of RL and RL* were adjusted to produce a titration where the weighted average peak in the final titration point (a magenta contour) is observed at approximately the same position as in Panel A. The equilibrium constant of the induced fit step, K, was set to 3 to favor RL*, and the reverse rate constant, k_2_, was set to 3000 s^-1^. This selection results in the exchange rate constant for the induced fit transition of *k*_*ex*_ *= k*_1_ + *k*_2_ *= Kk*_2_ + *k*_2_ *=* 12,000 *s*^−1^. Frequency differences between RL and RL* in Panel A are 0.15 ppm in proton and 1.5 ppm in nitrogen dimension corresponding to Δ*ω* of 565 s^-1^ in either dimension at a magnetic field of 14 Tesla. Comparing this Δ*ω* with k_ex_ of 12,000 s^-1^ makes us anticipate that an addition of the induced fit step to the fast-exchange U-5R model would make the titration appear in even faster exchange regimes than in Panel A. Indeed, not only the induced fit step was set in the fast exchange, but the frequency difference of the ligand binding step, *R* + *L* ↔ *RL*, was also reduced in this simulation (namely, the RL frequencies are closer to the R frequencies in Panel B than in panel A).

Yet, Figure 9.B reveals a dramatic change in the opposite direction. The “marching” fast-exchange weighted-average peak simulated for the U-5R mechanism in the Figure 9.A gave way to a much slower, intermediate exchange pattern despite the obvious fast-exchange regime in *every transition* in the U-5R-RL model. This phenomenon resembles an observation reported earlier for the three-state mechanisms in one-dimensional simulations^19^.

To identify the cause of such a non-intuitive exchange regime trend, we will reduce the U-5R-RL to its parent models and simulate the titration line shapes while retaining fast kinetic rate constants. First, we will remove all but one non-binding isomers of R arriving at the U-R-RL model, which includes both pre-existing equilibrium and the induced fit steps (Figure 10.A). For the ease of comparison, the R* frequency was picked to produce a weighted average resonance in the ligand-free simulation appearing at the same ppm position as in Figure 9.B (blue contour). The visual appearance of the exchange regime remained unchanged, which means that multiplicity of the non-binding conformations of the free receptor (R*) makes little contribution to the apparent deceleration of the exchange regime.

**Figure 10.**
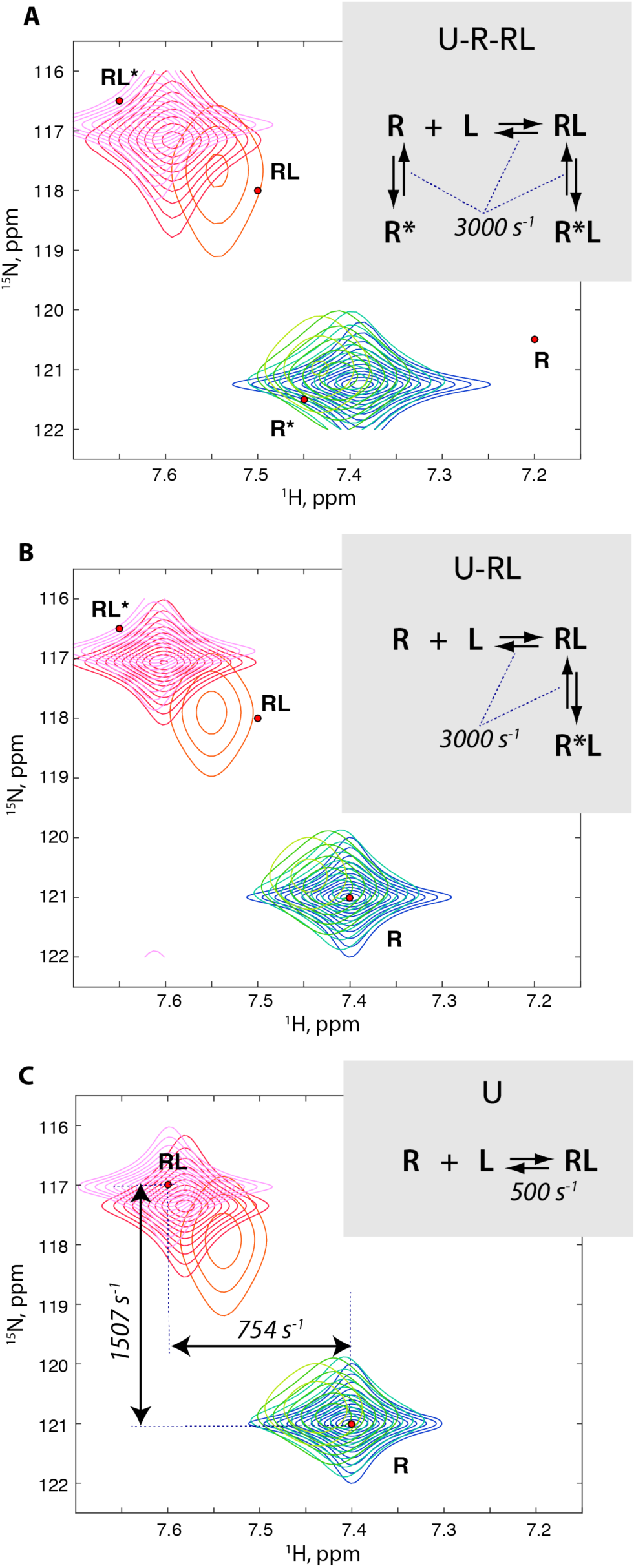
2D ^1^H-^15^N HSQC NMR spectra simulated for U-R-RL (**A**), U-RL (**B**), and U (**C**) models with all kinetic constants (k_2,A_ and k_2,B_) set to 3000 s^-1^ in **A** and **B** panels, while k_2,A_ was 500 s^-1^ in the **C** panel. Titration schedules and the color scheme is the same as in Figure 9. The chemical shifts of species in U-R-RL, U-RL, and U simulations were adjusted to have weighted-aver-age peaks appear at the same ppm positions as in the first and last titration points in Figure 9. For full details of the simulation setup see *simulations/U_R_RL, simulations/U_RL*, and *simulations/U_slow*.

Removing the pre-existing equilibrium step from U-R-RL produces a standard induced-fit model, U-RL (Figure 10.B). Comparison of its simulated spectral pattern with U-5R-RL (Figure 9.B) and U-R-RL (Figure 10.A) reveals that the exchange regime appears similarly decelerated despite the fast exchange in both steps of the mechanism. However, if the induced step is removed (making the basic two-state U model, Figure 10.C), such an intermediate exchange pattern may only be produced if the dissociation rate constant, k_2_, is reduced to as low as 500 s^-1^ (compare to the frequency differences between R and RL of 750 s^-1^ in ^1^H and 1500 s^-1^ in ^15^N dimensions in Figure 10.C). In addition, the three-state model with pre-existing equilibrium, U-R, made out of U-R-RL (Figure 10.A) by removing the induced fit step demonstrates the expected fast-exchange pattern shown in Supporting Figure 6. Therefore, the observed deceleration of the exchange regime in presence of the induced fit is not a mere consequence of addition of another kinetic step to the binding mechanism, instead, it appears to be a specific property imparted by the induced fit transition.

These observations lead us to two general conclusions. First, it is obvious that the exchange regime in the titration series for a multi-step mechanism cannot be easily ‘eye-balled’ by comparing the expected exchange rate constants with frequency differences. The full simulation of the titration series is required, and the general NMR line shape theory needs further development to explain fundamental reasons for the non-intuitive behaviors noted above. As the first step, it may be necessary to introduce additional terms such as *local* exchange regime (corresponding to a specific transition in the mechanism—based on a classical comparison of *k*_*ex*_ and Δ*ω*), and a *global* or *apparent* exchange regime (based on observed degree of broadening and the peak shift pattern in the titration series).

Second, we may anticipate that, in practical NMR titrations, the experimental systems involving induced fit transitions are likely to appear in a deceptively slower exchange regime while underlying reaction kinetics may be quite fast. Alternatively, observation of an apparent fast exchange regime in the course of a titration may be a sign of the molecular interaction mechanism as simple as a single-step binding (U), or the conformational selection model (U-R, …, U-5R, etc) without a significant contribution from the induced fit.

#### Multiplicity of binding-competent conformations in equilibrium with closed free and bound states

The 5U-R-RL model in Figure 3 represents a situation distinct from what we discussed in the previous sections—a mechanism where the major conformation of the receptor is not capable of interacting with ligand. Transient formation of the minor conformers (either through opening, unfolding, or other mechanisms) leads to productive interactions with the ligand. Yet the structural stability of these complexes may be low leading further to formation of a tightly bound state, which can no longer release a ligand (“locked”). For purposes of illustrating the spectral properties of the 5U-R-RL mechanism, we will simulate a case study while asking the following question: are the NMR line shapes in a titration sensitive to the *local* exchange regime in the C transitions (isomerizations among R isomers and among RL isomers)?

Figure 11.A demonstrates a titration series where I chose the chemical shifts of amide proton spins to be sensitive to opening-closing transitions (R* ⇒ R’,…, R’’’’’; and RL*, …, RL***** ⇒ RL) while nitrogen-15 spins reported on the ligand binding. The reverse rate constants in *all* transitions are set to 1 s^-1^, and equilibrium isomerization constants are chosen to create an approximately equal split between R* and all R isomers and between RL* and all RL isomers (see Supporting Figure 7 for the populations plot). Titration in Figure 11.A proceeds in a slow exchange regime revealing individual peaks of initial and final forms (R* and RL*) as well as peaks of all binding-competent isomers of R (R’, R’’, …., R’’’’’) and the first-encounter complexes (RL’, RL’’...,..., RL’’’’’).

**Figure 11.**
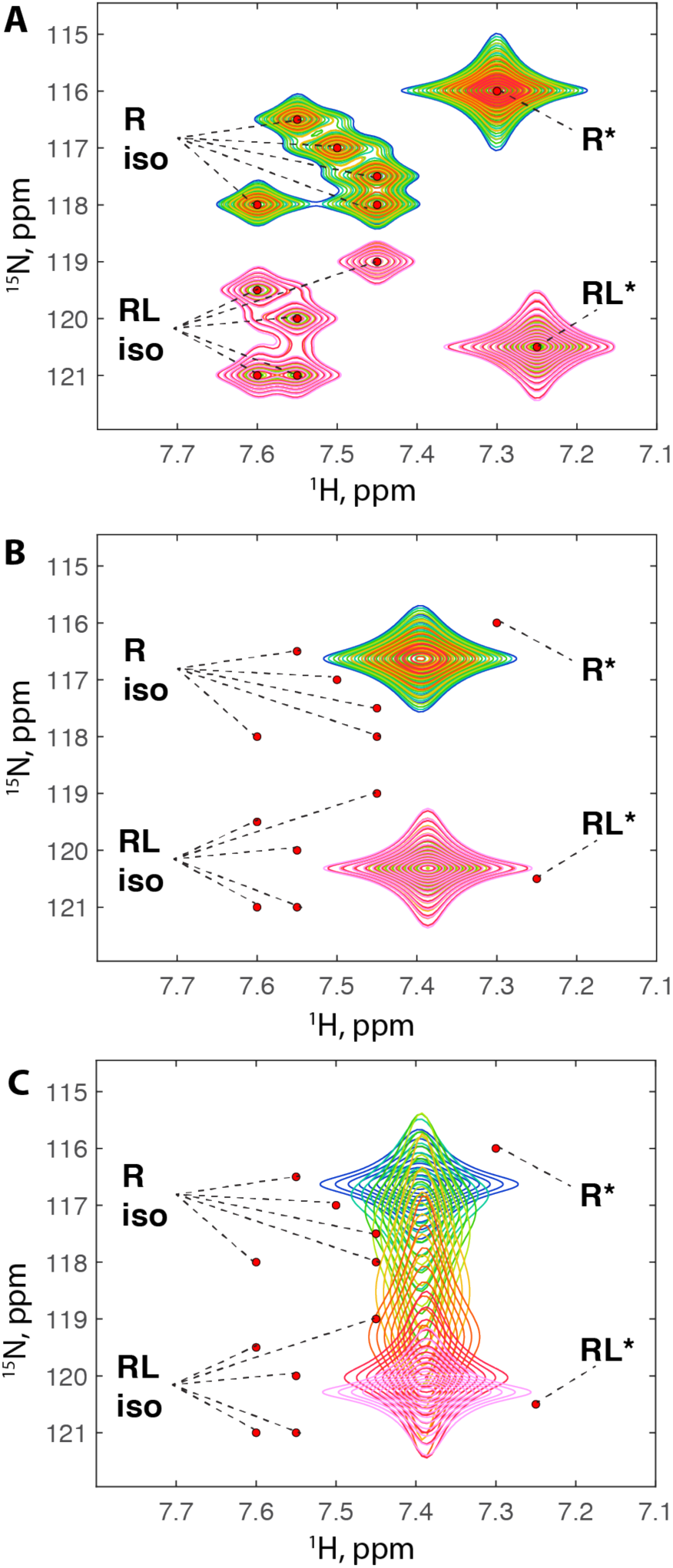
2D ^1^H-^15^N HSQC NMR spectra simulated for 5U-R-RL models. Dissociation constant for ligand binding is 10^−6^. Reverse rate constants for transitions A, B, and C were set as (A) k_2,A_= k_2,B_ =k_2,C_ =1 s^-1^, (B) k_2,A_=1 s^-1^, k_2,B_ = 3000 s^-1^, k_2,C_ =1 s^-1^, and (C) k_2,A_=3000 s^-1^, k_2,B_ = 3000 s^-1^, k_2,C_ =1 s^-1^. Titration schedules and the color scheme is the same as in Figure 9. For full details of the simulation setup see *simulations/5U_R_RL_individual*.

Figure 11.B shows the simulation for the same setup with isomerization in B transitions (R* ⇒ R’,…, R’’’’’; and RL*,..., RL***** ⇒ RL) accelerated towards the fast exchange limit. We are no longer able to see peaks of individual isomers of R and RL. Despite the distinct chemical shifts of all R isomers, the fast exchange with a common R* species results in a single homogeneously broadened weighted average peak. The same is observed for RL isomers. Clearly, the extremely slow exchange regime in C transitions is not detectable if the opening/closing transitions are fast. We may further modify the simulation setup in Figure 11.C by increasing ligand binding kinetics to 3000 s^-1^. Again, the titration series appears in the fast-intermediate exchange regime leaving no hint of underlying slow kinetics of the mutual isomerization transitions within the R and RL isomer families. Based on these observations, it might be expected that acceleration of kinetics in C transitions will produce no easily detectable features in the titration line shapes—confirmed by simulations in the Supporting Figure 8.

#### Summary

This paper presented analysis of spectral simulations for ligand-binding mechanisms involving multiple isomerization transitions. Two-dimensional simulations of explicit mathematical models revealed that exchange regimes in coupled steps of complex mechanisms interact in a non-intuitive manner. NMR practitioners may utilize such simulations to anticipate appearance of the experimental NMR spectra based on specific hypotheses about underlying molecular mechanisms. The next most interesting application of such analysis would be in its reverse form: starting from experimental data to quantitatively resolve details of complex mechanisms operating at the molecular level. This task, however, is complicated by the sheer number of fitting parameters one will have to determine for multi-state models. One obvious way to reduce chances of overfitting the data is to perform a simultaneous fit of line shape titrations for many resonances in sets of spectra recorded for different nuclei and at several different temperatures. Resonances from multiple spins at different molecular sites will display a certain chemical shift variation thus (hopefully) appearing in distinct exchange regimes, while the temperature variation will alter exchange regimes directly. Global fitting of such datasets combined with data from complementary biophysical techniques could enable detalization of biomolecular mechanisms with un-paralleled accuracy.

## Acknowledgement

The author is grateful to Rafael Brüschweiler, Katherine Henzler-Wildman, Blake Hill, Patrick Loria, and Jeffrey Peng for helpful discussions.

## Supporting Information

**Supporting Figure 1.**
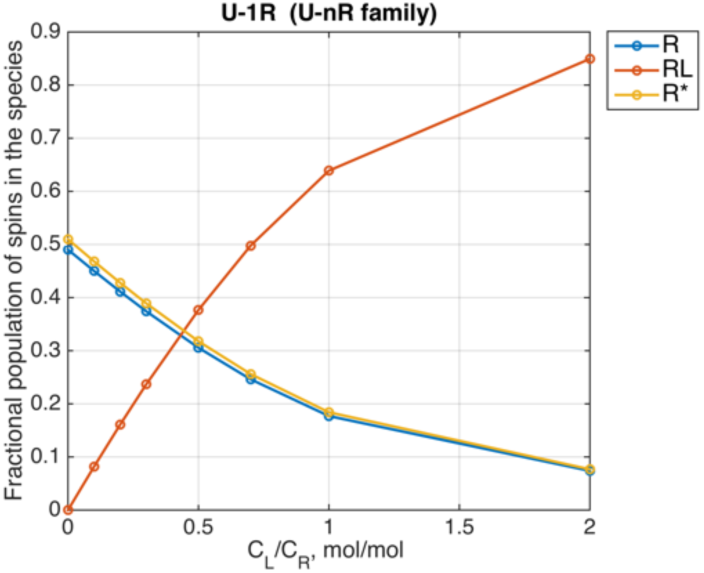
Populations of species calculated for the course of a titration of the U-1R system with the binding affinity constant of K_A_= 10^5^ (K_d_=10^−5^) and the isomerization constant K_B_=1.04. The K_B_ value was chosen to be equal to 1.04 instead of 1.0 for visual clarity: to make the population curves for R and R* distinct on the plot yet create nearly equal populations of the isomers. The initial concentration of the receptor was set to 0.1 mM.

**Supporting Figure 2.**
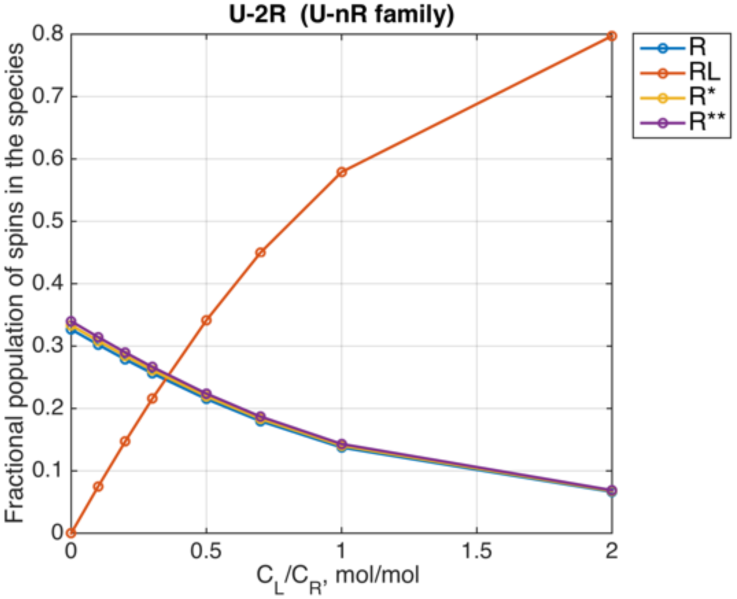
Populations of species calculated for the course of a titration of the U-2R system. The binding affinity constant was 10^5^ (K_d_=10^−5^), and the isomerization constants were set to 1.02 and 1.04 to assure nearly equal population of the isomers yet create visual distinction of the curves on the plot. The initial concentration of the receptor was set to 0.1 mM. For full details of the simulation setup see *simulations/ U_2R_Bint_Afast_3*.

**Supporting Figure 3.**
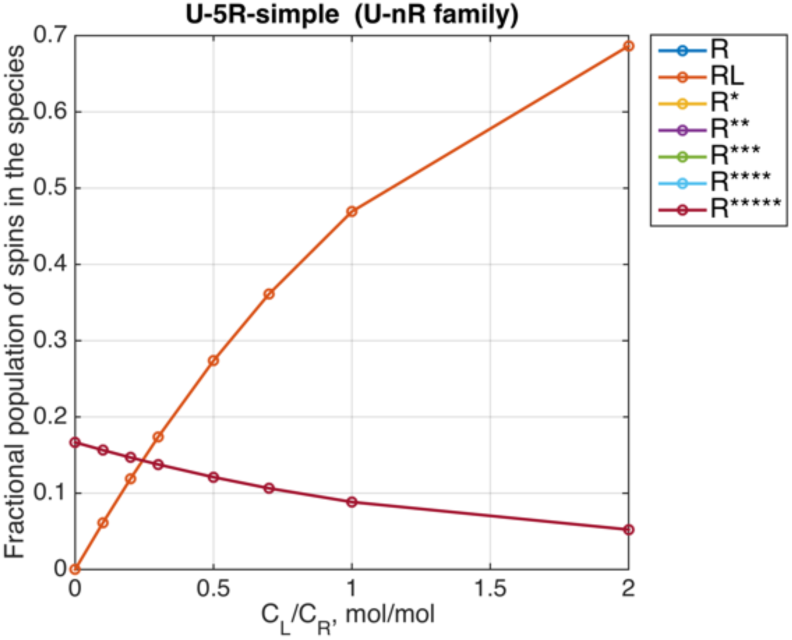
Populations of species calculated for the course of a titration of the U-5R system with the binding affinity constant of K_A_=10^5^, the isomerization constants set to 1 for all five of isomerization transitions. The initial concentration of the receptor was 0.1 mM. For full details of the simulation setup see *simulations/ U_5R_transient_narrowing*.

**Supporting Figure 4.**
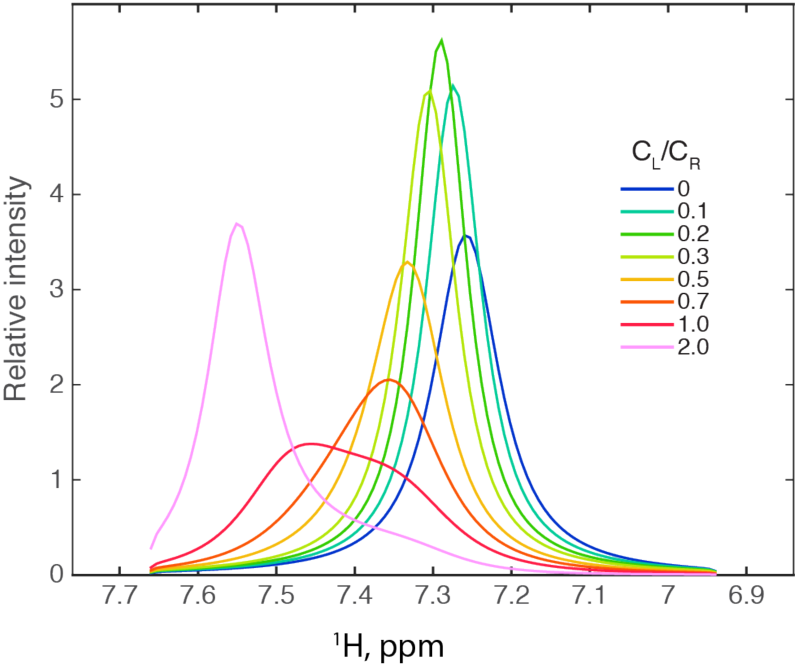
The projection of the two-dimensional spectra onto the proton dimension for the U-5R model simulation in Figure 8. The one-dimensional traces were obtained by integration of the nitrogen dimension of the corresponding 2D HSQC spectra.

**Supporting Figure 5.**
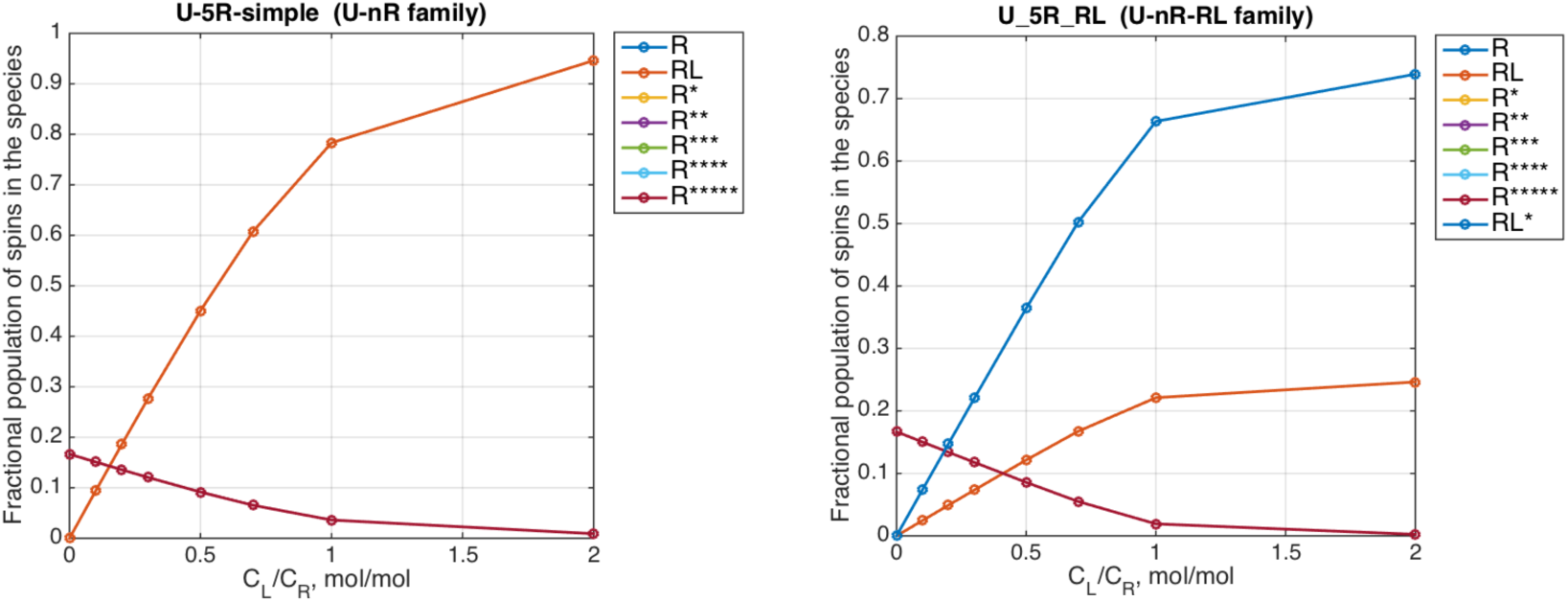
Populations of species calculated for the course of a titration of the U-5R and U-5R-RL systems in Figure 9. The initial concentration of the receptor was 0.1 mM. For full details of the simulation setup see *simulations/U_5R* and *simulations/U_5R_RL*.

**Supporting Figure 6.**
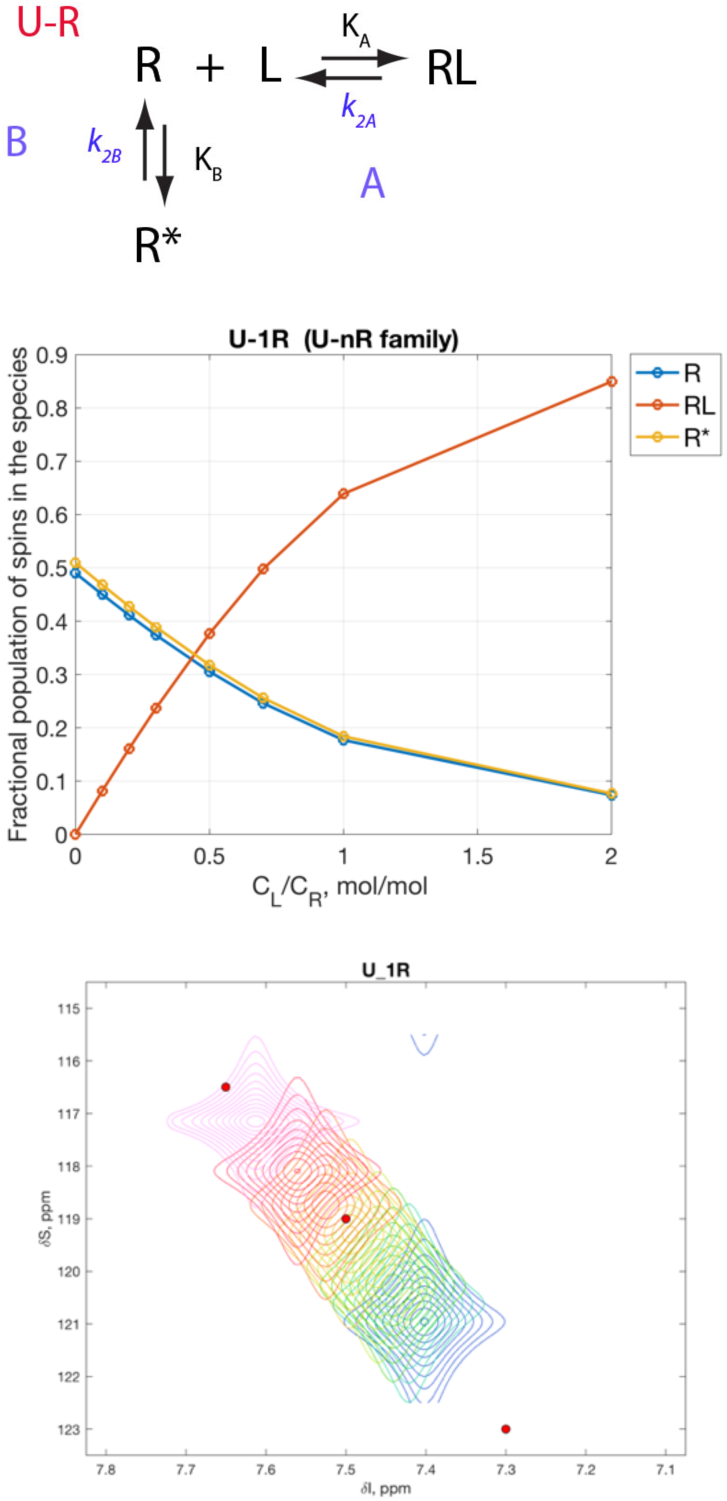
Simulations for U-1R model (top panel): populations of species (middle panel); 2D ^1^H-^15^N HSQC NMR spectra (bottom panel). Kinetic constants (k_2,A_ and k_2,B_) set to 3000 s^-1^. Full details of simulations are found in a setup file *simulations/U_1R_Bfast_Afast*.

**Supporting Figure 7.**
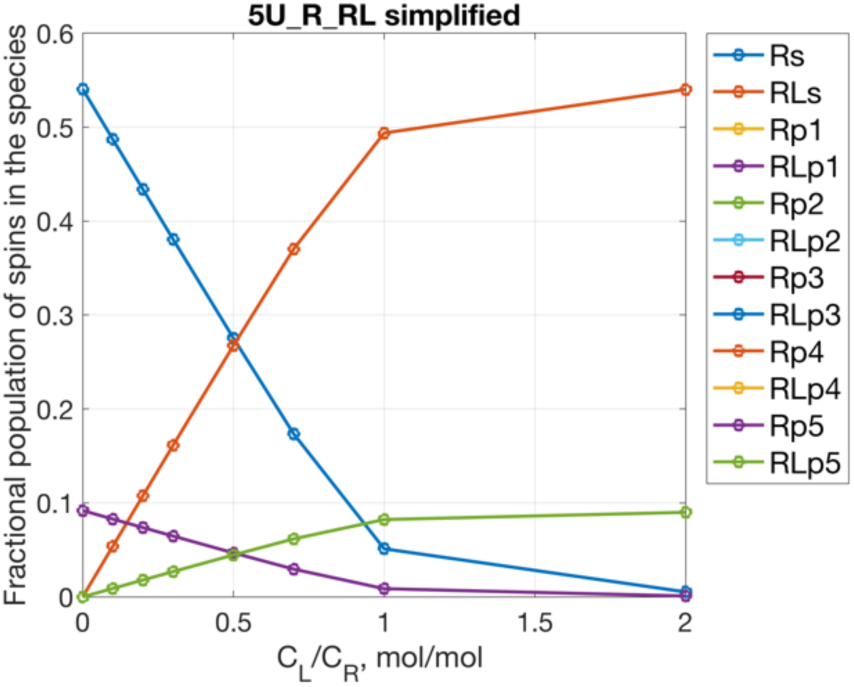
Populations of species calculated for the course of a titration of the 5U-R-RL systems in Figure 11. The initial concentration of the receptor was 0.1 mM. Populations of R-primed species are all the same because equilibrium constants in C transitions were set to 1, therefore, their curves are overlapped (Rp1, …, Rp5). Populations of RL-primed species are also identical for the same reason (labeled RLp1, …, RLp5). For full details of the simulation setup see *simulations/5U_R_RL.Cfast*.

**Supporting Figure 8.**
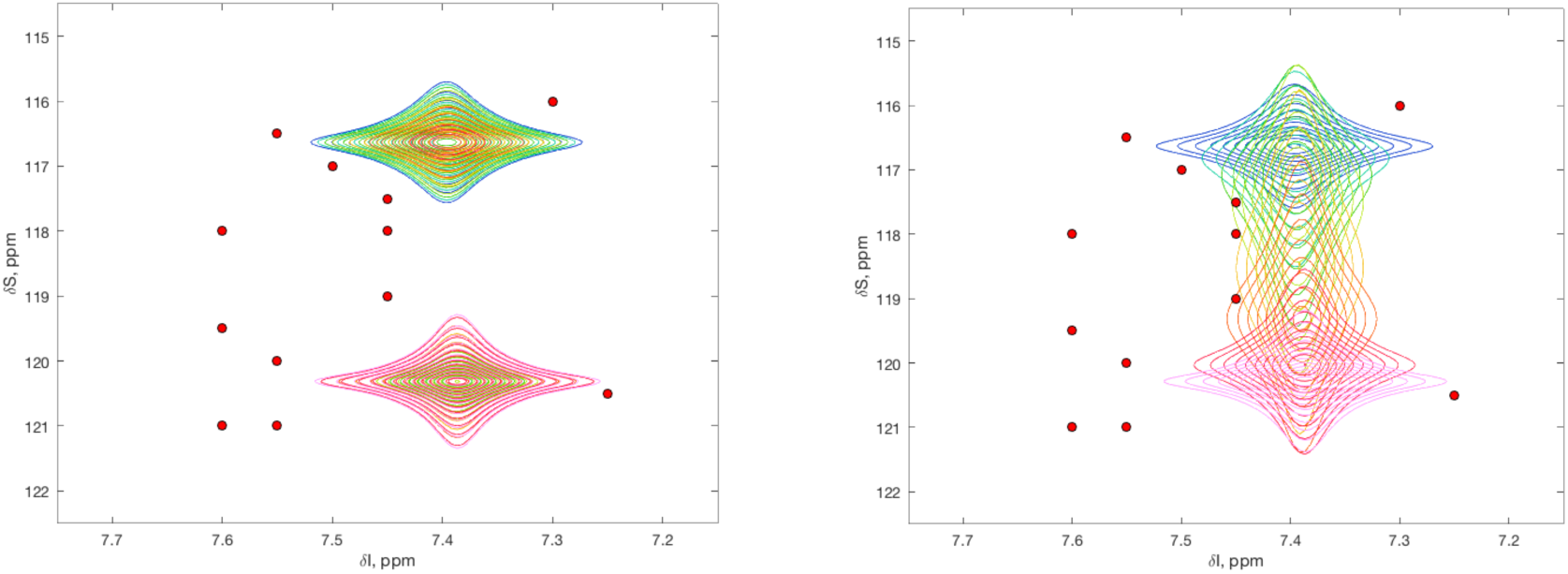
2D ^1^H-^15^N HSQC NMR spectra simulated for 5U-R-RL model with the reverse rate constants for C transitions set to 3000 s^-1^. All other parameters in the left and right panels are the same as in Figure 11.B and Figure 11.C, respectively. For full details of the simulation setup see *simulations/5U_R_RL.Cfast/As_Bf_Cf* and *simulations/5U_R_RL.Cfast/Af_Bf_Cf*.

## References

1. Rogers, J.M., Oleinikovas, V., Shammas, S.L., Wong, C.T., De Sancho, D., Baker, C.M. & Clarke, J. Interplay between partner and ligand facilitates the folding and binding of an intrinsically disordered protein. Proceedings of the National Academy of Sciences of the United States of America 111, 15420–15425 (2014).

2. Arai, M., Sugase, K., Dyson, H.J. & Wright, P.E. Conformational propensities of intrinsically disordered proteins influence the mechanism of binding and folding. Proceedings of the National Academy of Sciences of the United States of America 112, 9614–9619 (2015).

3. Das, T. & Eliezer, D. Membrane interactions of intrinsically disordered proteins: The example of alpha-synuclein. Biochimica Et Biophysica Acta-Proteins and Proteomics 1867, 879–889 (2019).

4. Koshland, D.E. The key-lock theory and the induced fit theory. Angewandte Chemie International Edition Engl. 33, 2375–2378 (1994).

5. Liu, Z.J., Jiang, L., Li, W.Z., Han, Y.Z. & Lai, L.H. Conformational changes accompanying with the binding of protein and ligand. Acta Chimica Sinica 58, 772–776 (2000).

6. Daniels, K.G., Tonthat, N.K., McClure, D.R., Chang, Y.C., Liu, X., Schumacher, M.A., Fierke, C.A., Schmidler, S.C. & Oas, T.G. Ligand Concentration Regulates the Pathways of Coupled Protein Folding and Binding. Journal of the American Chemical Society 136, 822–825 (2014).

7. Hubbard, S.R. Autoinhibitory mechanisms in receptor tyrosine kinases. Frontiers in Bioscience 7, D330–D340 (2002).

8. Hansen, M.D.H. & Kwiatkowski, A.V. Control of Actin Dynamics by Allosteric Regulation if Actin Binding Proteins. in International Review of Cell and Molecular Biology, Vol 303, Vol. 303 (ed. Jeon, K.W.) 1–25 (2013).

9. Chuenchor, W., Jin, T., Ravilious, G. & Xiao, T.S. Structures of pattern recognition receptors reveal molecular mechanisms of autoinhibition, ligand recognition and oligomerization. Current Opinion in Immunology 26, 14–20 (2014).

10. McConnell, H. Reaction rates by nuclear magnetic resonance. J. Chem. Phys. 28, 430–431 (1958).

11. Rao, B.D.N. Nuclear magnetic resonance line-shape analysis and determination of exchange rates. in Methods in Enzymology 279–311 (Academic Press, 1989).

12. Kern, D., Kern, G., Scherer, G., Fischer, G. & Drakenberg, T. Kinetic analysis of cyclophilin-catalyzed prolyl cis/trans isomerization by dynamic NMR spectroscopy. Biochemistry 34, 13594–602 (1995).

13. Günther, U.L. & Schaffhausen, B. NMRKIN: Simulating line shapes from two-dimensional spectra of proteins upon ligand binding. Journal of Biomolecular NMR 22, 201–209 (2002).

14. Doucet, N., Khirich, G., Kovrigin, E.L. & Loria, J.P. Alteration of hydrogen bonding in the vicinity of histidine 48 disrupts millisecond motions in RNase A. Biochemistry 50, 1723–30 (2011).

15. De, S., Greenwood, A.I., Rogals, M.J., Kovrigin, E.L., Lu, K.P. & Nicholson, L.K. Complete Thermodynamic and Kinetic Characterization of the Isomer-Specific Interaction between Pin1-WW Domain and the Amyloid Precursor Protein Cytoplasmic Tail Phosphorylated at Thr668. Biochemistry 51, 8583–96 (2012).

16. Mittag, T., Schaffhausen, B. & Gunther, U.L. Direct observation of protein-ligand interaction kinetics. Biochemistry 42, 11128–11136 (2003).

17. Mittag, T., Schaffhausen, B. & Gunther, U.L. Tracing kinetic intermediates during ligand binding. Journal of the American Chemical Society 126, 9017–9023 (2004).

18. Mittag, T., Franzoni, L., Cavazzini, D., Schaffhausen, B., Rossi, G.L. & Gunther, U.L. Retinol modulates sitespecific mobility of apo-cellular retinol-binding protein to promote ligand binding. Journal of the American Chemical Society 128, 9844–9848 (2006).

19. Kovrigin, E.L. NMR line shapes and multi-state binding equilibria. J Biomol NMR 53, 257–270 (2012).

20. Waudby, C.A., Ramos, A., Cabrita, L.D. & Christodoulou, J. Two-dimensional NMR lineshape analysis. Scientific Reports 6, 8 (2016).

21. Shinya, S., Ghinet, M.G., Brzezinski, R., Furuita, K., Kojima, C., Shah, S., Kovrigin, E.L. & Fukamizo, T. NMR line shape analysis of a multi-state ligand binding mechanism in chitosanase. J Biomol NMR 67, 309–319 (2017).

22. Kaplan, J.I. & Fraenkel, G. NMR of Chemically Exchanging Systems, (Academic Press, 1980).

23. Waudby, C.A., Frenkiel, T. & Christodoulou, J. Cross-Peaks in Simple Two-Dimensional NMR Experiments from Chemical Exchange of Transverse Magnetisation. Angewandte Chemie-International Edition 58, 8784-8788(2019).

24. Palmer, A.G., 3rd. Chemical exchange in biomacromolecules: past, present, and future. J Magn Reson 241, 3–17 (2014).

